# Use of FFPE-Derived DNA in Next Generation Sequencing: DNA extraction methods

**DOI:** 10.1101/521534

**Authors:** Samantha J. McDonough, Aditya Bhagwate, Zhifu Sun, Chen Wang, Michael Zschunke, Joshua A. Gorman, Karla J. Kopp, Julie M. Cunningham

**Affiliations:** Medical Genome Facility, Mayo Clinic, Rochester MN, U.S.A.; Department of Health Sciences Research, Division of Biomedical Statistics and Informatics, Mayo Clinic, Rochester MN, U.S.A.; Pathology Research Core, Mayo Clinic, Rochester MN, U.S.A.; Department of Laboratory Medicine & Pathology, Mayo Clinic, Rochester, MN, U.S.A

## Abstract

Archival tissues represent a rich resource for clinical genomic studies, particularly when coupled with comprehensive medical records. Use of these in NGS is a priority. Nine formalin-fixed paraffin-embedded (FFPE) DNA extraction methods were evaluated using twelve FFPE samples of varying tissue types. Quality assessment included total yield, percent ds DNA, fragment analysis and multiplex PCR. After assessment, three tissue types from four FFPE DNA methods were selected for NGS downstream evaluation, whole exome (WES) and targeted sequencing. In addition, two low input library protocols were evaluated for WES. Analysis revealed average coverage across the target regions for WES was ~20-30X for all four FFPE DNA methods. For the targeted panels, the highest molecular tag coverage was obtained with the Kingfisher FFPE extraction method. The genotype concordance was 99% for the commonly called variant positions between all four extraction methods with the targeted PCR NGS panel and 96% with WES.

## Introduction

Next generation sequencing (NGS) is rapidly becoming established in the clinic, predominantly in oncology but also as a means of diagnosis in individuals with unresolved medical issues. Archival tissue represents a singularly valuable resource for disease oriented research, particularly when combined with comprehensive medical records such as that of the Mayo Clinic. However, DNA extracted from such samples can vary widely in quality due to age, fixation conditions, DNA-protein crosslinking, and inhibitors, which may impact downstream genomic analyses. Samples are typically obtained in the operating room so how they are handled, as well as time exposed to formalin both contribute to potential DNA damage but are typically outside the control of investigators[1].

With the use of highly sensitive NGS applications it is imperative that the FFPE DNA extractions used in these assays be of the best quality obtainable. Variation in pre-processing may lead to inconsistencies in detection of mutations or viral presence[2,3] and variation in both quantity and quality of DNA extracted at four commercial laboratories was reported by Arreaza et al[4]. Thus, not only DNA quality, but the methods used to extract DNA contribute to performance in downstream assays.

Approaches for extraction of DNA from FFPE samples have been compared by others[2,5,6,7,8,9,10]; one used NGS for downstream analysis for two samples only[7], while Bonnet et al[5] used whole exome sequencing NGS on 42 samples, comparing three DNA extraction approaches but not comparing the same samples extracted by the three methods. Schweiger et al[11], using a controlled fixation of 72 hours for the FFPE-DNA, noted equivalence of three methods but no data were shown; QIAamp was used for seven FFPE samples and revealed that copy number and mutation analyses were possible. Other NGS studies of FFPE-derived DNA extracted by a single method have been reported[11, 12, 13, 14, 15, 16, 17, 18]. Kerick et al varied input for a targeted capture-based from 500ng-1.5ug of FFPE-DNA of one of five patients and showed comparable performance using standard library preparation[17].

Recent developments of FFPE DNA extraction processes include: new deparaffinization solutions, repair strategies, and magnetic bead technology. These improvements along with the need to profile archival tissues led us to evaluate several DNA extraction methods, assess low input library preparations for such samples, and use the preferred methods in whole exome and targeted panel sequencing NGS applications. In addition, we applied DNA quality metrics to assess the degree of fragmentation of FFPE DNA. DNA was extracted from sequential sections of twelve blocks of paraffin-embedded, formalin fixed from several tissue types using nine commercially available extraction methods, including both manual and automated processes. These samples were selected to represent a spectrum of quality, from highly cellular to those with high adipose tissue content that typically yield poorer quality DNA. After DNA quality and quantity were assessed, four of the processes were chosen to be evaluated for performance in NGS technologies. These included two library preparation protocols for low input DNA whole exome sequencing (WES), as limited DNA yields from FFPE samples are common, and two targeted DNA panels.

## Material and Methods

### Samples

Following an approved IRB protocol, twelve paraffin blocks were selected from Mayo’s Pathology Research Core laboratory’s Control and Assay Development Paraffin Preserved Tissue archive. Tissues included normal and diseased sections from breast, colon, lung, pancreas, along with two unique normal tonsil tissues and two different sections of brain tissue (brain stem and cerebellum). Ten micrometer sections were sequentially cut from each block using a standard microtome (Leica Rotary Microtome RM2235, Leica Biosystems, Buffalo Grove, IL) and placed on slides. Slides were scraped and material was placed into a 1.5 ml microcentrifuge tube for DNA extraction.

### Manual and Automated FFPE Extractions

We compared both manual and automated extraction methods, evaluated extraction time, throughput and quality. Manual protocols evaluated for the study were KAPA Express Extract kit (KAPA Biosystems, Wilmington, MA, USA), Promega Reliaprep^TM^ FFPE gDNA Miniprep system (Promega, Fitchburg, WI, USA), QIAGEN QIAamp^®^ FFPE tissue kit, and QIAGEN GeneRead DNA FFPE kit (QIAGEN, Germantown, MD, USA). Automated extraction methods evaluated for the study were QIAGEN QIAsymphony DNA mini kit (QIAsymphony SP), QIAGEN GeneRead DNA FFPE kit (QIAcube), Maxwell RSC DNA FFPE Kit (Promega Maxwell^®^ RSC), PerkinElmer chemagic FFPE DNA kit (chemagic MSM 1; Perkin Elmer, Baesweiler, Germany), and Applied Biosystem’s MagMAX^™^ FFPE DNA Isolation Kit (Applied Biosystems, Foster City, CA, USA) (KingFisher Duo; Thermo-Fisher Scientific, Waltham MA, U.S.A.). All extractions were performed using one 10 μM section from each of the 12 tissue blocks. All sections were deparaffinized using QIAGEN’s Deparaffinization Solution following manufacturer’s guidelines, exceptions were QIAamp extraction protocol which used xylene and KAPA Express Extract which contains its own extraction buffer and enzyme. Manual and automated protocols for each method were followed according to manufacturers’ guidelines. The only modification made was to the QIAamp FFPE tissue protocol which used an overnight lysis incubation time of 56°C instead of the suggested 1 hour.

### Quality

DNA quality for each extracted sample was measured by evaluating quantity, purity, amount of double stranded DNA, and fragment length. Samples were quantified using Qubit^™^ dsDNA HS or BR Assays (ThermoFisher Scientific, Waltham, MA, USA) and NanoDrop^™^ NT-1000 spectrophotometer readings; the purity and amount of ds DNA was calculated using the ratio of Qubit to Nanodrop readings. Fragment length and degradation were assessed using the Advanced Analytical Fragment Analyzer^™^ High Sensitivity Large Fragment Analysis kit (Advance Analytical, Ankeny, IA, USA) and a multiplex PCR assay (Life Science Innovations, Qualitative Multiplex PCR Assay for Assessing DNA Quality from FFPE Tissues, and Other sources of Damaged DNA Issue 23, SigmaAldrich, St. Louis, MO) which uses amplicon size to determine degradation and fragment size.

### Sequencing

FFPE DNA samples and one CEPH control (Centre d’Etude du Polymorphisme Humain, Coreill Institute, Camden, NJ, USA) were subjected to two different low input library preparations for whole exome sequencing (WES), NEBNext^®^ Ultra II DNA Library Prep (New England BioLabs Inc. Ipswich, MA, USA) and ThruPLEX^®^ DNA-seq Kit (Rubicon Genomics, Ann Arbor, MI, USA). Library preparations were made following manufacturer’s guidelines using an input of 50ng of DNA. Library concentrations were assessed and approximately 500ng of each sample library was enriched using Agilent’s SureSelect XT Target Enrichment System V5+UTR (Agilent, Santa Clara, CA, USA). Libraries were quantified and sequenced thirteen per lane, PE 150bp on an Illumina HiSeq 4000 (Illumina, San Diego, CA, USA). To further investigate the influence of DNA FFPE extraction methods on downstream next generation sequencing applications, this same set of thirteen samples was prepared following manufacturer’s guidelines using the QIAGEN QIAseq^™^ Targeted Human Comprehensive Cancer Panel with an input of 40ng FFPE DNA from each sample. All libraries were sequenced in one lane per panel on an Illumina HiSeq 4000, PE 150bp.

### Analysis

Bioinformatics analysis for the WES was performed using in-house DNA analysis workflow (GenomeGPS v4.0.1). The reads were first aligned to the GRCh37 build of the human reference genome using BWA-MEM with the default parameter settings. After alignment, the reads were re-aligned and re-calibrated using Genome Analysis Toolkit (GATK) Indel Realigner to optimize the mapping around indels. Variant calling was then performed on the realigned reads using GATK Haplotype Caller and the called variants were functionally annotated using an in-house developed genomics annotation tool BioR. The QIAseq panel analysis was performed using QIAGEN’s NGS data analysis portal. The portal performs appropriate read trimming, generating consensus reads using unique molecular indexes (UMIs), and variant calling using QIAGEN’s barcode-aware variant caller “smCounter”, followed by variant annotation. We used a cutoff of 5x for concordance analyses.

## Results

### DNA Extraction

The methods used to extract DNA from the 12 FFPE samples are shown in Table 1 along with the acronym used for each in this report; modifications were not added to the manufacturers’ protocols with one exception noted above.

**Table 1.** DNA Extraction methods.

| FFPE DNA Extract Kit | Process | Abbreviation |
| --- | --- | --- |
| KAPA Express Extract Kit | Manual | KE-M |
| Promega Reliaprep<sup>™</sup> FFPE gDNA Miniprep System | Manual | PR-M |
| QIAGEN GeneRead<sup>™</sup> DNA FFPE Kit | Manual | QGR-M |
| QIAGEN QIAamp<sup>®</sup> DNA Mini Kit (Mayo's Biospecimens Accessioning and Processing Lab protocol) | Manual | QA-M |
| QIAGEN QIAasympohy DSP DNA Mini Kit | Automated | QS-A |
| QIAGEN QIAcube GeneRead<sup>™</sup> DNA FFPE Kit | Automated | QGR-A |
| Promega Maxwell<sup>®</sup> RSC DNA FFPE Kit | Automated | PM-A |
| PerkinElmer Chemagic MSM1 FFPE DNA Kit | Automated | PEC-A |
| ThermoFisher KingFisher<sup>™</sup> Duo MagMAX<sup>™</sup> FFPE DNA Isolation Kit | Automated | TKM-A |

All of the methods used a proteinase (Proteinase K in all but KE-M which uses a thermostable protease NOS). Three of the four manual methods used silica based mini-elute columns to bind DNA; KE-M did not include a clean-up step. QGR-M adds a repair step that removes formalin crosslinks and de-aminated cytosines. The automated methods all use magnetic beads to isolate the DNA, with the exception of the automated GeneRead protocol which employs the QIAcube (silica based) for column purification. KE-M offered the shortest workflow compared the others, but its lack of a clean-up step outweighed the rapid protocol. The five automated protocols offered the most ease of use, with TKM-A being the most fully automated protocol.

### DNA Quality Assessment

Data for DNA yield and dsDNA content for the twelve different tissue types used on the four manual and five automated FFPE DNA extraction methods are shown in Fig 1A-C. Among the tissues, DNA extracted from brain stem and normal breast tissues had the lowest yields of DNA, tumors yielded more DNA than their normal counterparts, and tissue from tonsils yielded the highest (Fig 1A). Among the manual methods, QA-M and QGR-M methods were better performing when yield was considered, while QGR-A and TKM-A were the better among automated methods (Fig1B). When considering percentage of dsDNA, the template required for most downstream assays, QA-M, QGR (both manual and automated), QS-A and TKM-A yielded higher dsDNA percentages (Fig 1C). 260/280 ratios were above 1.8 for all methods except KE-M (S1).

**Fig 1.**
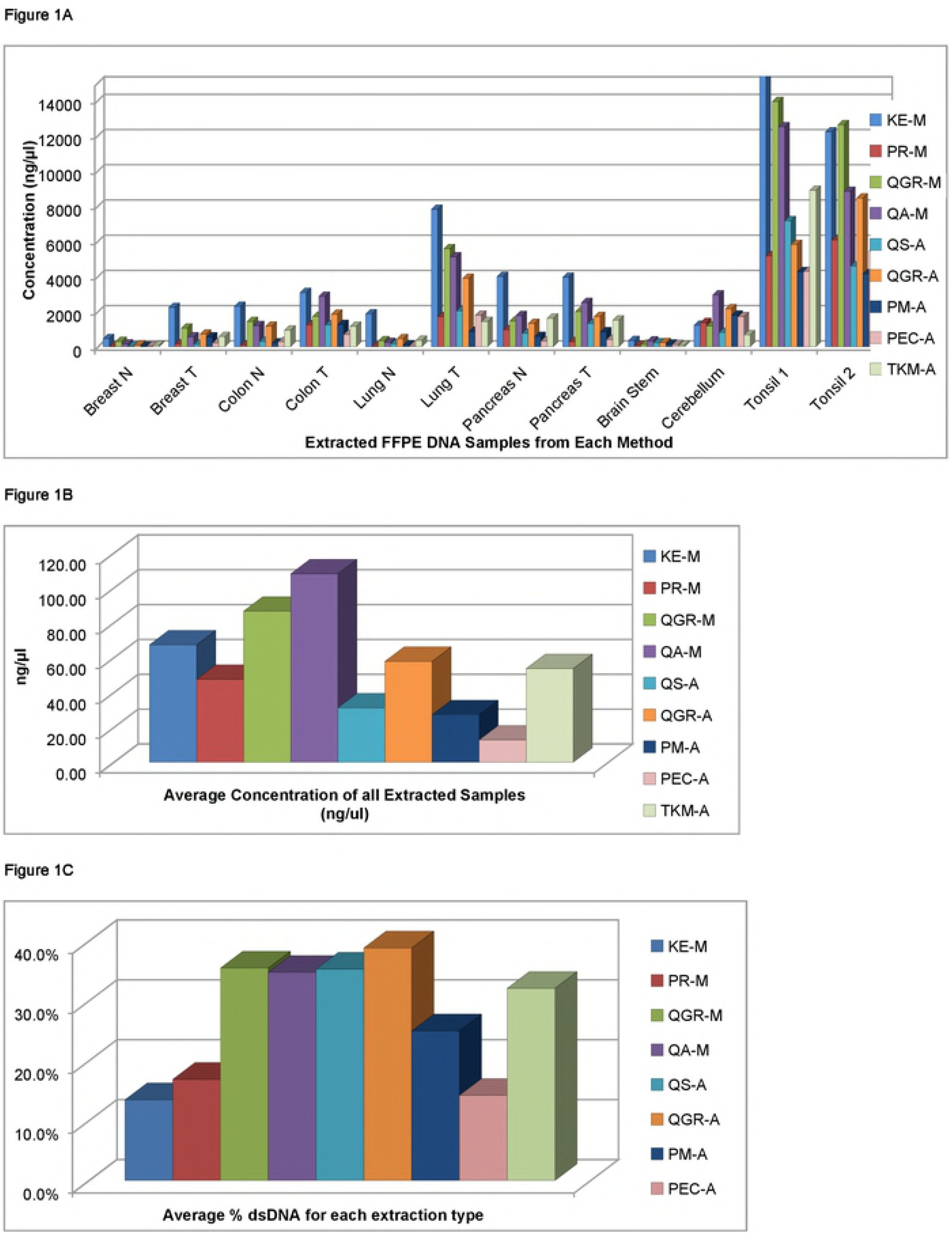
DNA quality metrics. Each extraction method is designated by a specific colour (see key) which is maintained in all figures except Fig 5. A: Concentration for each sample of extracted DNA, B: Average concentration for all samples extracted, C: Percent double stranded DNA for all samples.

Fragment analysis data are shown in Fig 2A-C with the full data set in S2.

**Fig 2.**
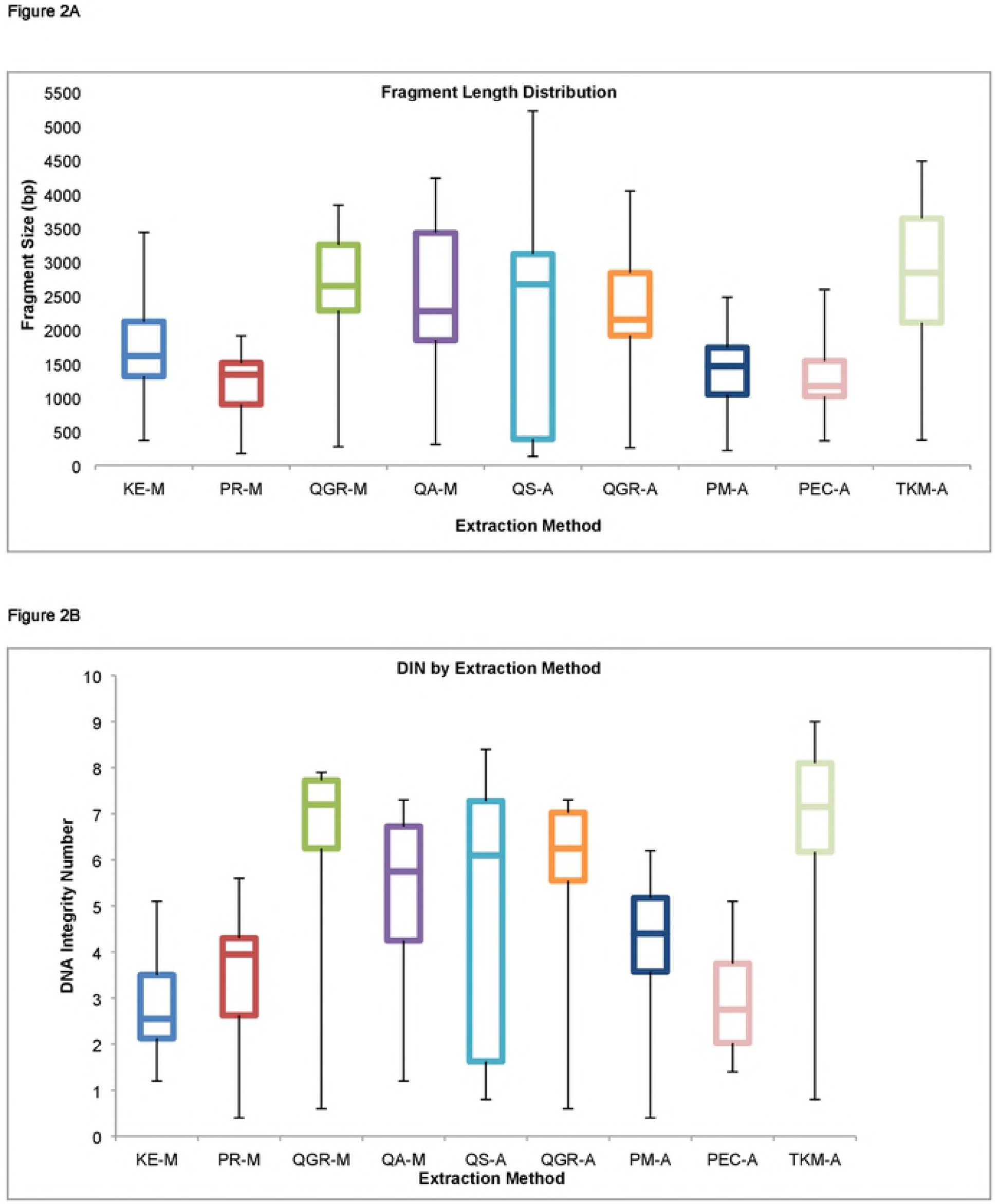
Fragment Analysis. A: Median DNA fragment lengths with 75-25 percentiles for each method, B: Median DNA Integrity numbers (DIN) for each method; a size threshold of 500bp was selected for assessment. C: Percent of fragments <200bp, 201-1,000bp, 1,001-20,000bp for three representative tissue types.

The median fragment lengths of each of the nine approaches are shown in Fig 2A. Several methods were poor overall at preserving fragment length, notably KE-M, PR-M, PM-A and PEC-A, while QS-A was more variable. Those that do preserve fragment length most consistently were QGR (manual and automated), QA-M, PM-A and TKM-A methods. Figure 2B shows the DNA Integrity Number which reflects the findings for fragment length. Figure 2C shows the percentage of DNA from three samples (breast tumor, pancreas tumor and cerebellum) of varying quality at 1-200bp, 201-1000pb and 1001-20,000bp for four methods. More fragmented samples such as brain stem and cerebellum have most DNA <1000bp by most methods. However, for the tumor samples, QGR (manual and automated), QA-M, PM-A and TKM-A methods yielded larger percentages of higher molecular weight DNA. Three of these methods, QIAGEN GeneRead (GR-A or GR-M), Promega Maxwell (PM-A) and QIAGEN QIAamp (QA-M or QA-A), were evaluated in Bonnet et al[5]; QA and GR were less fragmented than PM-A, having longer median fragment lengths, in agreement with findings in this report.

Fragment analysis gives a sense of the fragmentation of the DNA; however whether it is effective in downstream applications may not follow. Thus, we evaluated performance in a multiplex PCR to determine how well amplicons of different size are amplified compared to a control sample. Data from the multiplex PCR are shown in S3. Results are variable based on method and tissue type. KE-M generally produced poorer results, likely due to the absence of a DNA clean up step. Poorer quality DNA (brain stem, cerebellum) generally produced <20 percent of amplifiable amplicons above 132-196 base pairs with most of the methods. Of all the methods, PEC-A, QS-A, and KE-M produced poor quality results while QGR (both manual and automated) and TKM-A results show consistent amplification of larger amplicons among the samples.

DNA extracted by QGR-M, QA-M, QGR-A and TKM-A methods were chosen to evaluate in downstream NGS applications based on yield, percent dsDNA and fragment length. Three of the twelve tissue types evaluated were chosen for this NGS assessment based on their quality scores, ranging from severe to moderate degradation; cerebellum, breast tumor and pancreas tumor.

### Next Generation Sequencing

For WES, two low input library preparation methods were used along with the four DNA extraction methods described previously; coverage metrics, read duplication rates and fragment insert sizes for each DNA extraction and library preparation method combination are shown in S4. DNA from all methods provided good mean median coverage (23.7X), except for the highly degraded cerebellum samples, with Ultra II providing slightly better coverage (25.88X) than ThruPlex (21.5X). Additionally, the distribution of coverage across the targeted bases as shown in Fig 3A and S5 further suggests that the UltraII method is able to cover some targeted bases even at 75X, while the Thruplex coverage distribution generally tapers off beyond 50X. A circos plot of raw coverage between methods for a subset of exons targeted is shown in Fig 3B.

**Fig 3.**
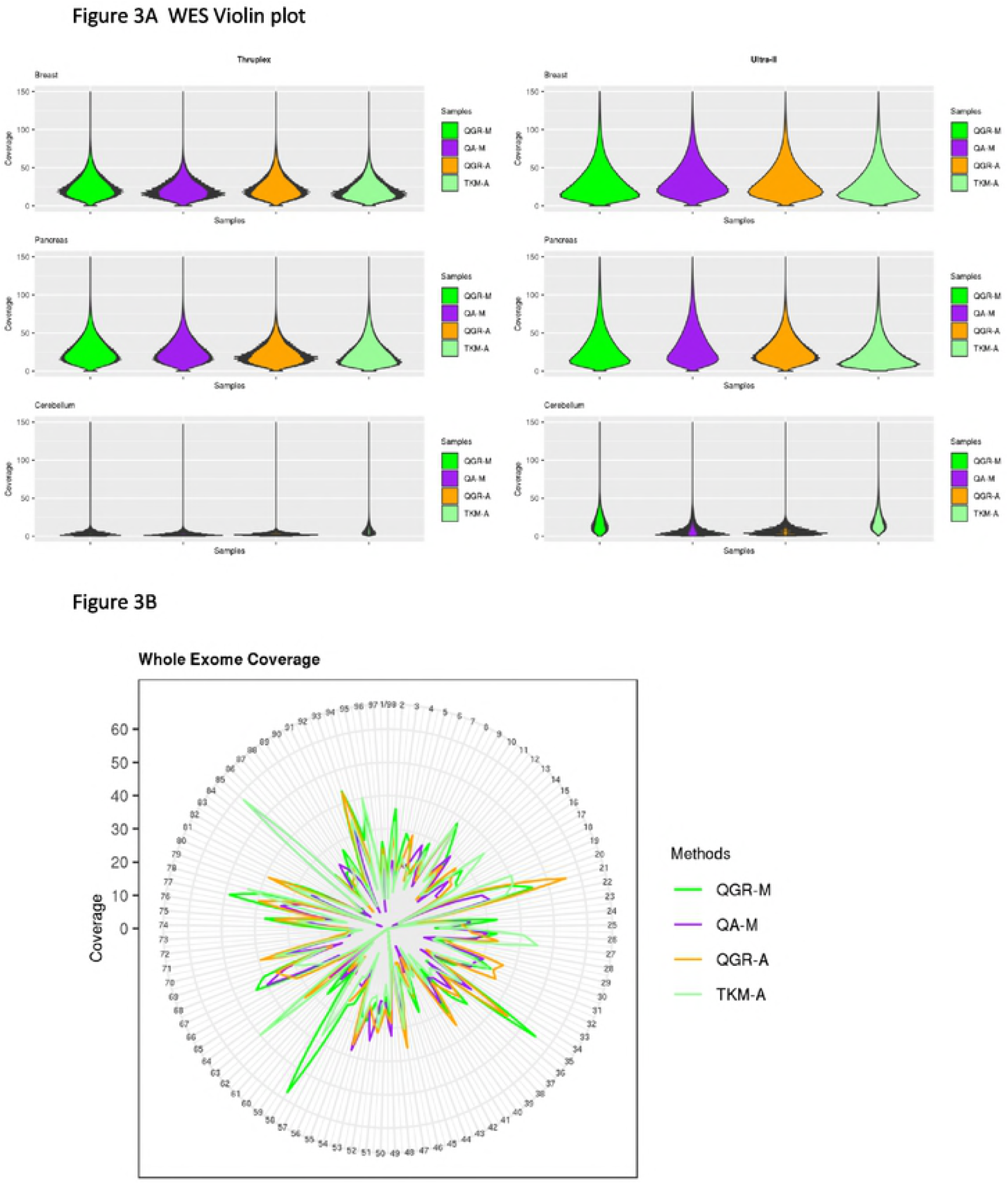
Raw coverage distribution for the four DNA extraction methods. A) Across all targeted bases as seen in the Thruplex (left panel) and Ultra-II (right panel) methods and B) Across a subset of the targeted regions with the Ultra II method and the Breast samples. The overall coverage is very similar for the four methods. The cerebellum samples show significantly lower coverage compared to other tissues due to the highly degraded nature of the samples. The CIRCOS plot the coverage lines within are noted on the left (0X the inner most to 60X the outer most concentric lines).

Unlike Bonnet et al[5] who observed an approximate average median coverage difference of 13.5X between extraction methods for FFPE samples, we observed that both GR-A/M and QAM methods produced similar metrics between the four extraction methods with respect to coverage (Δ_coverage_ = 2.55X), percentage of bases with ≥ 20x coverage (Δ_pctbases ≥ 20X_ = 5.05 percent), percent bases mapping in target (Δ_target_ = 4.25 percent) and outside target regions (Δ_offtarget_ = 3.12 percent). However, we used a low DNA input (50ng) rather than 200ng used by Bonnet et al.

While we did not observe significant differences in duplication rates between the four DNA extraction methods (Δ_duplication_= 3.05 percent), we observed that the read duplication rates were lower with the Ultra II library preparation method (11.9 percent) than ThruPlex (26.21 percent). Also, for the most degraded sample, cerebellum, duplication rates were generally higher than for the other tissues types (S4).

For variant calling comparison between methods, we used genotype concordance as a measure to assess the reproducibility of called variants along with the number of variants detected with each method. Concordance was called between methods when the variant position and the variant genotype were identical between methods. Table 2A shows the number of variants called with each method, while Table 2B shows the genotype concordance between each pair of methods. An average genotype concordance of 96% for called variants was observed between the methods and alternate allele frequencies showed an average Pearson’s Score correlation of r^2^ 0.95.

**Table 2.** Number of Variants Called (A) and Percent Genotype Concordance (B)

| **Tissue** | **Breast** |  | **Pancreas** |  | **Cerebellum** |  |
| --- | --- | --- | --- | --- | --- | --- |
| **Methods** | TP | U-II | TP | U-II | TP | U-II |
| **QGR-M** | 70,463 | 70,523 | 70,462 | 70,951 | 55,316 | 71,807 |
| **QA-M** | 69,516 | 70,326 | 69,547 | 69,488 | 47,097 | 61,337 |
| **QGR-A** | 69,952 | 71,249 | 69,468 | 70,791 | 50,478 | 59,878 |
| **TKM-A** | 70,013 | 72,112 | 71,235 | 70,968 | 76,539 | 83,721 |

\*TP=ThruPlex; U-II= Ultra II

B42
| **Method** | **QGR-M** |  | **QGR-A** |  | **TKM-A** |  |
| --- | --- | --- | --- | --- | --- | --- |
|  | TP | U-II | TP | U-II | TP | U-II |
| **QA-M** | 95.56 | 96.70 | 95.60 | 96.72 | 95.43 | 96.62 |
| **QGR-M** | 100 | 100 | 95.77 | 96.59 | 95.45 | 95.41 |
| **QGR-A** | 100 | 100 | 100 | 100 | 95.58 | 96.55 |

\*TP=ThruPlex; U-II= Ultra II

For targeted sequencing, we evaluated performance in QIAGEN QIAseq Human Comprehensive Cancer (DHS-3501Z) and Human Breast Cancer (DHS-001Z) Panels. Coverage and insert size metrics for each sample are shown in S6. For the Comprehensive Cancer Panel, the highest average molecular tag (MT) coverage of 820.91X was obtained with the TKM-A, while the QGR-M, QA-M and QGR-A methods produced average MT coverages of 650.96X, 599.68X and 733.77X respectively. Similar coverage results were observed with the Breast Cancer Panel, with the TKM-A method producing the highest average molecular tag coverage of 1609.23X, and the QGR-M, QA-M and QGR-A methods resulting in 749.12X, 1089.31X and 731.02X respectively. Figure 4 Tables 3A and 3B show the number of variants called with the Comprehensive Cancer Panel and the genotype concordance between the variant calls for the four methods respectively. The average genotype concordance between the variant calls was found to be high at 99.59% between the four extraction methods.

**Fig 4.**
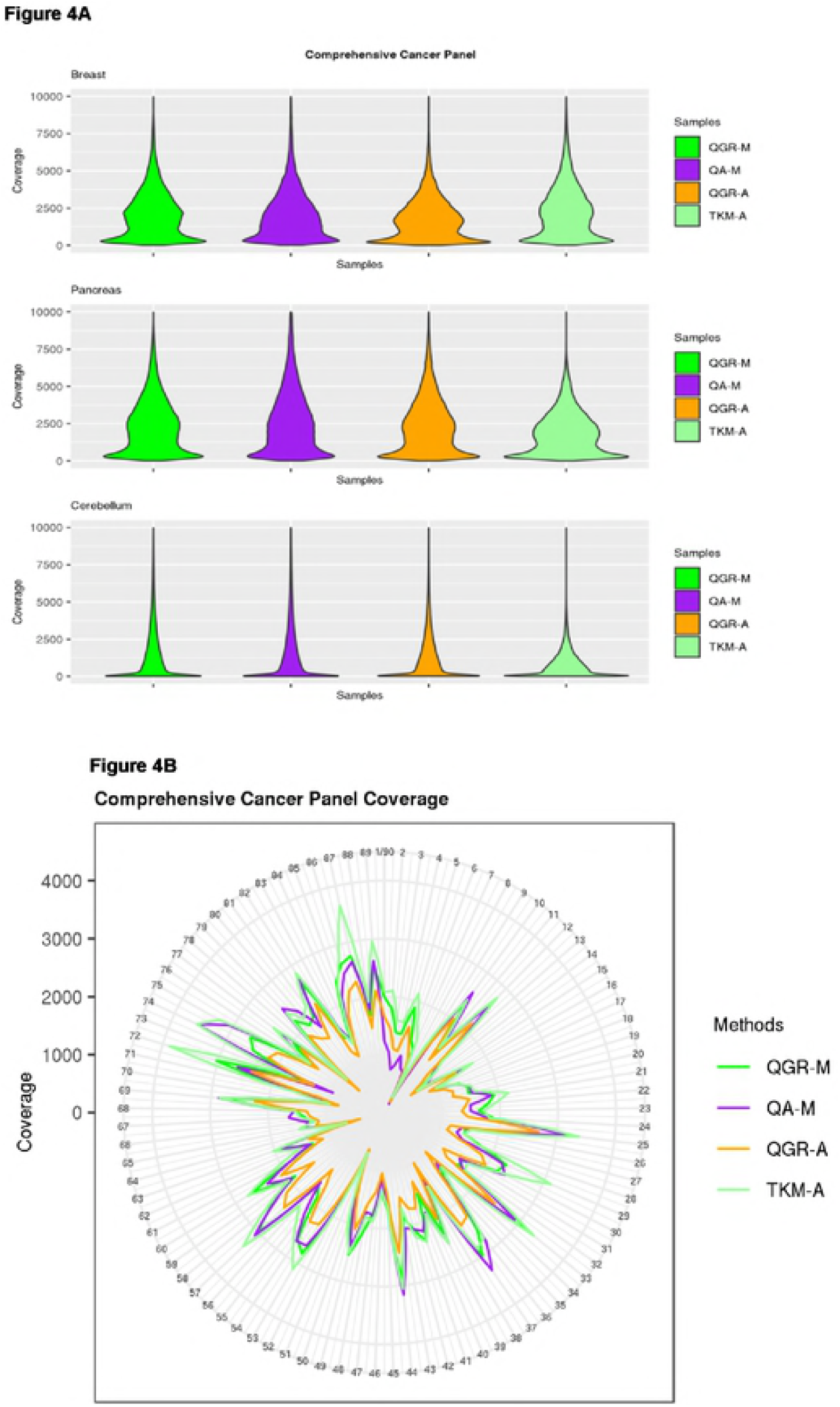
Raw Coverage distribution for the four DNA extraction methods. A) Across all targeted bases as seen with the Comprehensive Cancer Panel and B) Across a subset of the targeted regions in the Comprehensive Cancer Panel. The CIRCOS plot represents the regions targeted; the coverage lines within are noted on the left (0X the inner most to 4000X the outer most concentric lines).

**Table 3.**
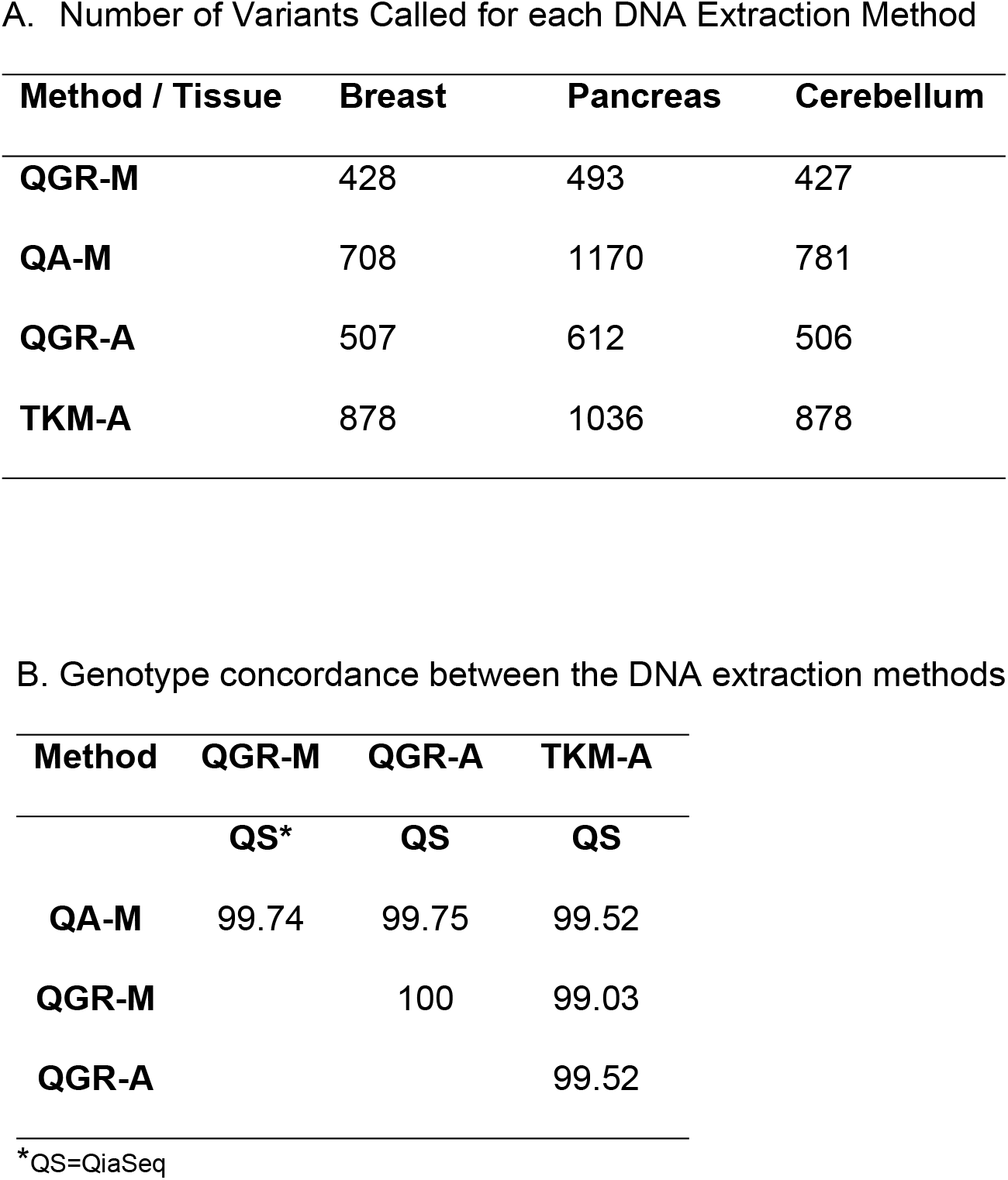
A. Number of Variants Called for each DNA Extraction Method

| Method / Tissue | Breast | Pancreas | Cerebellum |
| --- | --- | --- | --- |
| **QGR-M** | 428 | 493 | 427 |
| **QA-M** | 708 | 1170 | 781 |
| **QGR-A** | 507 | 612 | 506 |
| **TKM-A** | 878 | 1036 | 878 |

B. Genotype concordance between the DNA extraction methods
| Method | QGR-M | QGR-A | TKM-A |
| --- | --- | --- | --- |
|  | **QS\*** | **QS** | **QS** |
| **QA-M** | 99.74 | 99.75 | 99.52 |
| **QGR-M** |  | 100 | 99.03 |
| **QGR-A** |  |  | 99.52 |

\*QS=QiaSeq

The distribution of coverage for the comprehensive cancer panel for three representative samples is shown in (Fig 4.) Each of the four methods yielded some proportion of target bases even at a raw coverage value as high as 5000X but the proportion of targeted bases covered above 5000X drops off significantly. For the highly degraded cerebellum sample, the raw coverage drops off beyond 2500X. Figure 4B represents the raw coverage of a subset of targeted regions across the four methods.

### Variant analysis and FFPE artifacts

The sequencing data showed high concordance of genotypes between all NGS applications, indicating that any of these four methods could offer optimal yield and quality. Where the sequencing results differ is in variant calling with the Q-GR showing fewer variants called due to the enzymatic repair step which removes artificially induced uracil in the DNA. These artefacts are known to occur in FFPE-derived DNA [1, 19, 20, 21], due in part to augmented deamination of cytosine or adenosine [1, 19, 20, 21, 22] and show up in sequencing as C>T/G>A or A>G/T>C variants. More recently, Einaga et al[23] and Prentice et al[24] also noted possible artefactual mutations in FFPE samples, highlighting the role of project design and bioinformatics analyses. We observed that the QA-M method called the highest percentage of C>T/G>A mutations at 38.06 percent, while the QGR methods which include the enzymatic repair step called 36.65 percent of variants as C>T/G>A (Table 4).

**Table 4:** WES summary metrics.

|  | QGR-M | QGR-A | QA-M | TKM-A |
| --- | --- | --- | --- | --- |
| DNA median Fragment size | 2644* | 2146 | 2270.5 | 2838 |
| Median % of positions with >20X | 47.65/ 61.67 | 40.96/ 61.00 | 44.04/ 63.70 | 43.67/ 52.75 |
| Median coverage | 19.5/ 24.5 | 17 / 23.75 | 18.5 / 25.75 | 18 / 20.75 |
| % duplicated reads | 18/9.6* | 19.6/14 | 22.2/14.4 | 19.2/9.6 |
| C.T>G.A rate for SNV<br>(% of C>T/G>A mutations) | 36.64 | 37.5 | 38.06 | 36.59 |

\*with/without cerebellum

When looking at the sequence context of the C>T transitions[18], we found 43% to occur in CpG dyads. Additionally, we observed that ~32% of variants were also called as T>C/G>A, which have also been reported as artefacts resulting from FFPE DNA[20]. Fig 5A further shows the distribution of variant signatures seen across the Breast FFPE sample in whole exome sequencing (data only shown for the UltraII method); Fig 5B shows the same with the Comprehensive Cancer Panel.

**Fig 5.**
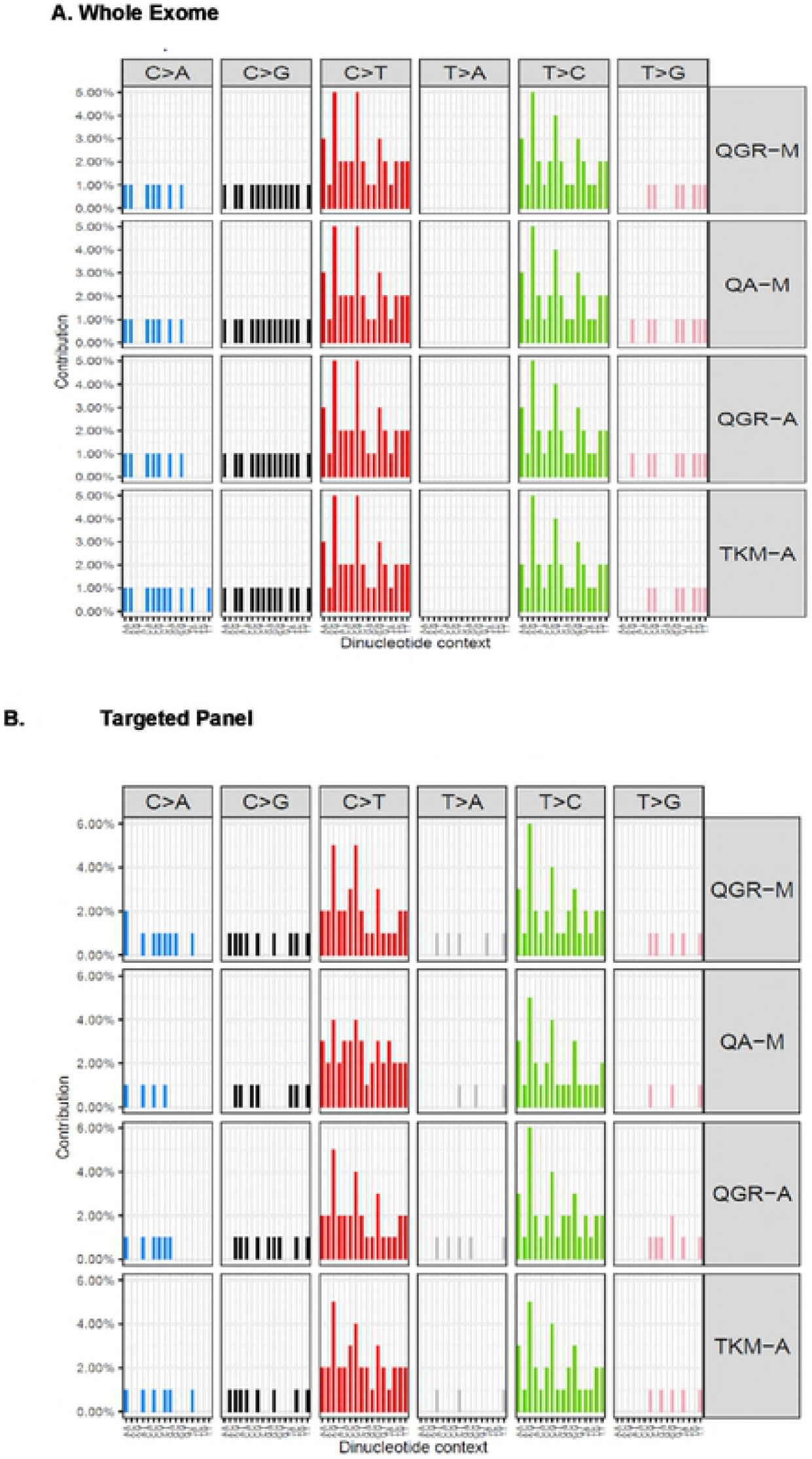
Signature of mutations found in the FFPE samples. A) Standard whole exome sequencing and B) Targeted Comprehensive Cancer Panel. A large proportion of the mutations are seen in the C>T context resulting from cytosine de-aminations in both datasets. The QGR method involving the repair step reduces but does not eliminate these C>T mutations.

## Discussion

DNA extraction methods for FFPE tissues vary in quality and quantity of resultant DNA, all of which may impact performance in downstream assays. In this report, we evaluated nine methods, including both manual and automated protocols, as the latter are preferred in to minimize potential errors in sample handling. QC evaluations were performed to determine the degree of DNA damage and which methods application might procure the best quality data; DNA from four extraction methods (manual and automated) were assayed for low input WES and amplicon based targeted sequencing based on yield, percent dsDNA and fragment length. For WES, the NEBNext Ultra II low input library preparation provided the lower duplicated reads, better coverage, and higher reads in capture regions than did those prepared using ThruPlex for the four extraction methods. The very fragmented DNA from cerebellum had lower reads in capture region and higher duplication rates for WES but was effectively profiled using an amplicon-based targeted approach.

The low input library preparation methods were very similar, the only difference being the type of bead used for the cleanup step, suggesting that small changes can impact downstream performance. There are new methods emerging for use of small inputs for NGS applications, which should further broaden the number of samples available for use.

There are some challenges using FFPE DNA in NGS. There were no substantial differences in sequencing results for the four selected protocols, neither was there any difference in the spectrum of variation found. The FFPE signature, of C>T transitions was similar in all tested methods. As in Spencer et al [18], the finding that C>T transitions often occurred in CpG dyads supports the observation that deamination of cytosine FFPE is a major source of artefactual variations in FFPE DNA[1]. This occurs in living cells and uracil-DNA glycosylase (UDG) removes the altered base; the abasic site is then restored to cytosine by base excision repair[1]. The GeneRead protocol includes a repair step (UDG) which ameliorates, but does not totally eliminate, these artefacts. Bioinformatics approaches to dealing with these artefacts will enhance the use of FFPE derived DNA in epidemiologic studies.

The strengths of this report are the evaluation of nine DNA extraction methods, including manual and automated, and evaluation of low input library preparation protocols. Sequential sections from a single block for each sample were used, to best assess each extraction approach. As the samples included in the study were anonymized, it was not possible to compare with matched frozen tissue samples.

## Supporting Information

**S1 Fig 1** Average 260/280 ratios of the 12 DNAs for each extraction method

**S2 Table 1** Percentage of fragments within each size range for all DNA extraction methods for the 12 samples.

**S3 Table 2** Multiplex PCR data expressed as percentage compared to a CEPH control. Highlighted columns represent those methods selected for evaluation in NGS.

**S4 Table 3** Whole Exome Sequencing Metrics for the two library preparation methods and four selected DNA extraction methods

**S5 Fig 2** Heat map showing the percentage of target bases covered by pancreas tumor, cerebellum and breast cancer samples, DNA extraction and library preparation methods.

**S6 Table 4** Metrics for the two targeted NGS panels

## Acknowledgements

This work was supported by the Mayo Clinic Center for Individualized Medicine. We would like to acknowledge the manifold contributions made by W. Edward Highsmith, who passed away 22^nd^ May 2018.

## Author contributions

### Conceptualization

Samantha McDonough, Julie Cunningham

### Laboratory processes

Samantha McDonough, Michael Zschunke, Joshua Gorman, Karla Kopp

### Bioinformatics analysis

Aditya Bhagwate, Chen Wang, Zhifu Sun

### Writing-original draft

Julie Cunningham, Samantha McDonough, Aditya Bhagwate.

## References

1. Do H, Dobrovic A (2015) Sequence artifacts in DNA from formalin-fixed tissues: causes and strategies for minimization. Clinical chemistry 61: 64–71.

2. Alvarez-Aldana A, Martinez JW, Sepulveda-Arias JC (2015) Comparison of five protocols to extract DNA from paraffin-embedded tissues for the detection of human papillomavirus. Pathology, research and practice 211: 150–155.

3. Kapp JR, Diss T, Spicer J, Gandy M, Schrijver I, et al. (2015) Variation in pre-PCR processing of FFPE samples leads to discrepancies in BRAF and EGFR mutation detection: a diagnostic RING trial. Journal of clinical pathology 68: 111–118.

4. Arreaza G, Qiu P, Pang L, Albright A, Hong LZ, et al. (2016) Pre-Analytical Considerations for Successful Next-Generation Sequencing (NGS): Challenges and Opportunities for Formalin-Fixed and Paraffin-Embedded Tumor Tissue (FFPE) Samples. International journal of molecular sciences 17.

5. Bonnet E, Moutet ML, Baulard C, Bacq-Daian D, Sandron F, et al. (2018) Performance comparison of three DNA extraction kits on human whole-exome data from formalin-fixed paraffin-embedded normal and tumor samples. PloS one 13: e0195471.

6. Dallol A, Al-Ali W, Al-Shaibani A, Al-Mulla F (2011) Analysis of DNA methylation in FFPE tissues using the MethyLight technology. Methods in molecular biology 724: 191–204.

7. Heydt C, Fassunke J, Kunstlinger H, Ihle MA, Konig K, et al. (2014) Comparison of pre-analytical FFPE sample preparation methods and their impact on massively parallel sequencing in routine diagnostics. PloS one 9: e104566.

8. Janecka A, Adamczyk A, Gasinska A (2015) Comparison of eight commercially available kits for DNA extraction from formalin-fixed paraffin-embedded tissues. Analytical biochemistry 476: 8–10.

9. Rabelo-Goncalves E, Roesler B, Guardia AC, Milan A, Hara N, et al. (2014) Evaluation of five DNA extraction methods for detection of H. pylori in formalin-fixed paraffin-embedded (FFPE) liver tissue from patients with hepatocellular carcinoma. Pathology, research and practice 210: 142–146.

10. Senguven B, Baris E, Oygur T, Berktas M (2014) Comparison of methods for the extraction of DNA from formalin-fixed, paraffin-embedded archival tissues. International journal of medical sciences 11: 494–499.

11. Schweiger MR, Kerick M, Timmermann B, Albrecht MW, Borodina T, et al. (2009) Genome-wide massively parallel sequencing of formaldehyde fixed-paraffin embedded (FFPE) tumor tissues for copy-number- and mutation-analysis. PloS one 4: e5548.

12. Astolfi A, Urbini M, Indio V, Nannini M, Genovese CG, et al. (2015) Whole exome sequencing (WES) on formalin-fixed, paraffin-embedded (FFPE) tumor tissue in gastrointestinal stromal tumors (GIST). BMC genomics 16: 892.

13. Bonfiglio S, Vanni I, Rossella V, Truini A, Lazarevic D, et al. (2016) Performance comparison of two commercial human whole-exome capture systems on formalin-fixed paraffin-embedded lung adenocarcinoma samples. BMC cancer 16: 692.

14. Carrick DM, Mehaffey MG, Sachs MC, Altekruse S, Camalier C, et al. (2015) Robustness of Next Generation Sequencing on Older Formalin-Fixed Paraffin-Embedded Tissue. PloS one 10: e0127353.

15. Johansson H, Isaksson M, Sorqvist EF, Roos F, Stenberg J, et al. (2011) Targeted resequencing of candidate genes using selector probes. Nucleic acids research 39: e8.

16. Van Allen EM, Wagle N, Stojanov P, Perrin DL, Cibulskis K, et al. (2014) Whole-exome sequencing and clinical interpretation of formalin-fixed, paraffin-embedded tumor samples to guide precision cancer medicine. Nature medicine 20: 682–688.

17. Kerick M, Isau M, Timmermann B, Sultmann H, Herwig R, et al. (2011) Targeted high throughput sequencing in clinical cancer settings: formaldehyde fixed-paraffin embedded (FFPE) tumor tissues, input amount and tumor heterogeneity. BMC medical genomics 4: 68.

18. Spencer DH, Sehn JK, Abel HJ, Watson MA, Pfeifer JD, et al. (2013) Comparison of clinical targeted next-generation sequence data from formalin-fixed and fresh-frozen tissue specimens. The Journal of molecular diagnostics : JMD 15: 623–633.

19. Hofreiter M, Serre D, Poinar HN, Kuch M, Paabo S (2001) Ancient DNA. Nature reviews Genetics 2: 353–359.

20. Marchetti A, Felicioni L, Buttitta F (2006) Assessing EGFR mutations. The New England journal of medicine 354: 526–528; author reply 526-528.

21. Solassol J, Ramos J, Crapez E, Saifi M, Mange A, et al. (2011) KRAS mutation detection in paired frozen and Formalin-Fixed Paraffin-Embedded (FFPE) colorectal cancer tissues. International journal of molecular sciences 12: 3191–3204.

22. Hofreiter M, Jaenicke V, Serre D, von Haeseler A, Paabo S (2001) DNA sequences from multiple amplifications reveal artifacts induced by cytosine deamination in ancient DNA. Nucleic acids research 29: 4793–4799.

23. Einaga N, Yoshida A, Noda H, Suemitsu M, Nakayama Y, et al. (2017) Assessment of the quality of DNA from various formalin-fixed paraffin-embedded (FFPE) tissues and the use of this DNA for next-generation sequencing (NGS) with no artifactual mutation. PloS one 12: e0176280.

24. Prentice LM, Miller RR, Knaggs J, Mazloomian A, Aguirre Hernandez R, et al. (2018) Formalin fixation increases deamination mutation signature but should not lead to false positive mutations in clinical practice. PloS one 13: e0196434.

